# The making of the oral microbiome in Agta hunter-gatherers

**DOI:** 10.1101/2022.05.03.490437

**Authors:** Begoña Dobon, Federico Musciotto, Alex Mira, Michael Greenacre, Rodolph Schlaepfer, Gabriela Aguileta, Leonora H. Astete, Marilyn Ngales, Vito Latora, Federico Battiston, Lucio Vinicius, Andrea B. Migliano, Jaume Bertranpetit

## Abstract

Ecological and genetic factors have influenced the composition of the human microbiome during our evolutionary history. We analyzed the oral microbiota of the Agta, a hunter-gatherer population where part of its members is adopting an agricultural diet. We show that age is the strongest factor modulating the microbiome, likely through immunosenescence as there is an increase of pathogenicity with age. Biological and cultural processes generate sexual dimorphism in the oral microbiome. A small subset of oral bacteria is influenced by the host genome, linking host collagen genes to bacterial biofilm formation. Our data also suggests that shifting from a fish/meat to a rice-rich diet transforms their microbiome, mirroring the Neolithic transition. All these factors have implications in the epidemiology of oral diseases. Thus, the human oral microbiome is multifactorial, and shaped by various ecological and social factors that modify the oral environment.

## Introduction

The composition and diversity of the human oral microbiota has been influenced by several factors during our evolutionary history^1,2^. Some are intrinsic biological characteristics of the host, such as age, sex, and genetic composition, while others such as diet, drinking water sources, oral hygiene, lifestyle and social interactions are external factors^2–4^. These factors modulate the physiological conditions of the oral cavity and affect the composition and diversity of the oral microbiota. While the oral microbiota is one of the most diverse sites in the human body and shows high variability between individuals, it remains stable within individuals over time^5^. Little is known about how the composition of the oral microbiome is modulated in populations adapted to the hunting and gathering niche, where the fully mature oral biofilm microbiome can be studied without the confounding effects of tooth brushing or professional dental cleaning, similar conditions to how the human oral microbiome must have evolved in the past^6^.

To investigate the multiple ecological and genetic factors shaping the human oral microbiome, we have analysed the oral microbiome of the Agta hunter-gatherers from the Philippines. The Agta are predominantly hunter-gatherers (fishing, hunting, and gathering)^7^, and while their main source of animal protein is obtained by riverine and marine spearfishing or by hunting, other activities such as inter-tidal foraging, wild food gathering, low-intensity cultivation, wage labour and trade complement their economy^8,9^. Interestingly, there is high variability among the Agta on the amount of hunting, gathering and sea foraging products that is traded for rice and other items (such as tobacco) with farming neighbours^7^, causing the Agta lifestyle to range from completely mobile foragers with a protein-rich diet, to settled low intensity farmers, with a rice-rich diet^7,9,10^.

To detect fine-scale variation in the oral microbiome of Agta hunter-gatherers, we collected saliva samples from 138 Agta, aged 5 to 65 years, sequencing the 16S rRNA region, and identified 5430 amplicon sequence variants (ASVs)^11^ belonging to 110 genera. To study the genetic host factors associated with the microbiome composition we also genotyped all individuals with the Axiom Genome-wide Human Origins array. We combined this information with additional individual data on household composition, age, sex, and diet, measured as the proportion of meals including meat/fish (any animal protein), and proportion of meals that consisted of only agricultural products (rice) (Supplementary Figure 1). By using this rich dataset, we have been able to discern the different contributions of age, sex, diet, and host genetics in the making of the oral environment.

## Results

### Factors influencing Agta oral microbiome composition

The Agta oral microbiome is mostly composed by *Firmicutes* (mean ASV prevalence = 33.1%, sd = 11.4), *Proteobacteria* (27±14.6%), *Actinobacteria* (15.5±8.3%) and *Bacteroidetes* (14.5±7.7%) (Supplementary Figure 2). To compare and identify the main ecological and social factors contributing to microbiome variation, we performed a constrained logratio analysis (LRA)^12^ on the bacterial genus abundance. Marginally, age explains 7.2% of the total logratio variance (P < 0.0001, based on 9999 permutations), sex explains 2.2% (P = 0.018), and diet 3.6% (P = 0.015). Altogether they explain 13.0% of the total logratio variance. We also applied a bipartite stochastic block model (biSBM) approach^13^ at the ASV level, where we assigned each bacterium to an individual, and then clustered the individuals according to the bacteria they have in common. We restricted the analysis to the Core Measurable Microbiota (CMM), that we define as ASVs present in at least 10% of the Agta to reduce random errors due to low-prevalent taxa (Supplementary Figure 3). The best model produced two clusters of people and three clusters of bacteria (Figure 1a). While we did not find differences in diet or proportions of sexes between the clusters of individuals (Supplementary Figure 4), they strongly differ in their age distribution: adults (mean age = 38 years old) and youth (mean age = 18 years old) (Welch t-test, t = 5.78, df = 71.35, P < 0.0001), with 55.48% of ASVs in the CMM being more associated with one of the clusters of individuals. Thus, while age, diet, and sex influence the composition of Agta microbiome, the biSBM singles out age as the main modulator of the hunter-gatherer core oral microbiome.

### Old age is associated with increased frequency of oral pathogens

To investigate the independent effects of ageing on the oral microbiome, we performed a LRA constrained with age and sex after partialing out the effects of diet. The resulting ordination shows that the effects of age and sex are mostly independent, with only few genera being affected by both variables (Supplementary Figure 5): as expected, the first dimension is associated with the age of the individuals, while the second dimension separates them according to sex (Figure 1b).

**Figure 1.**
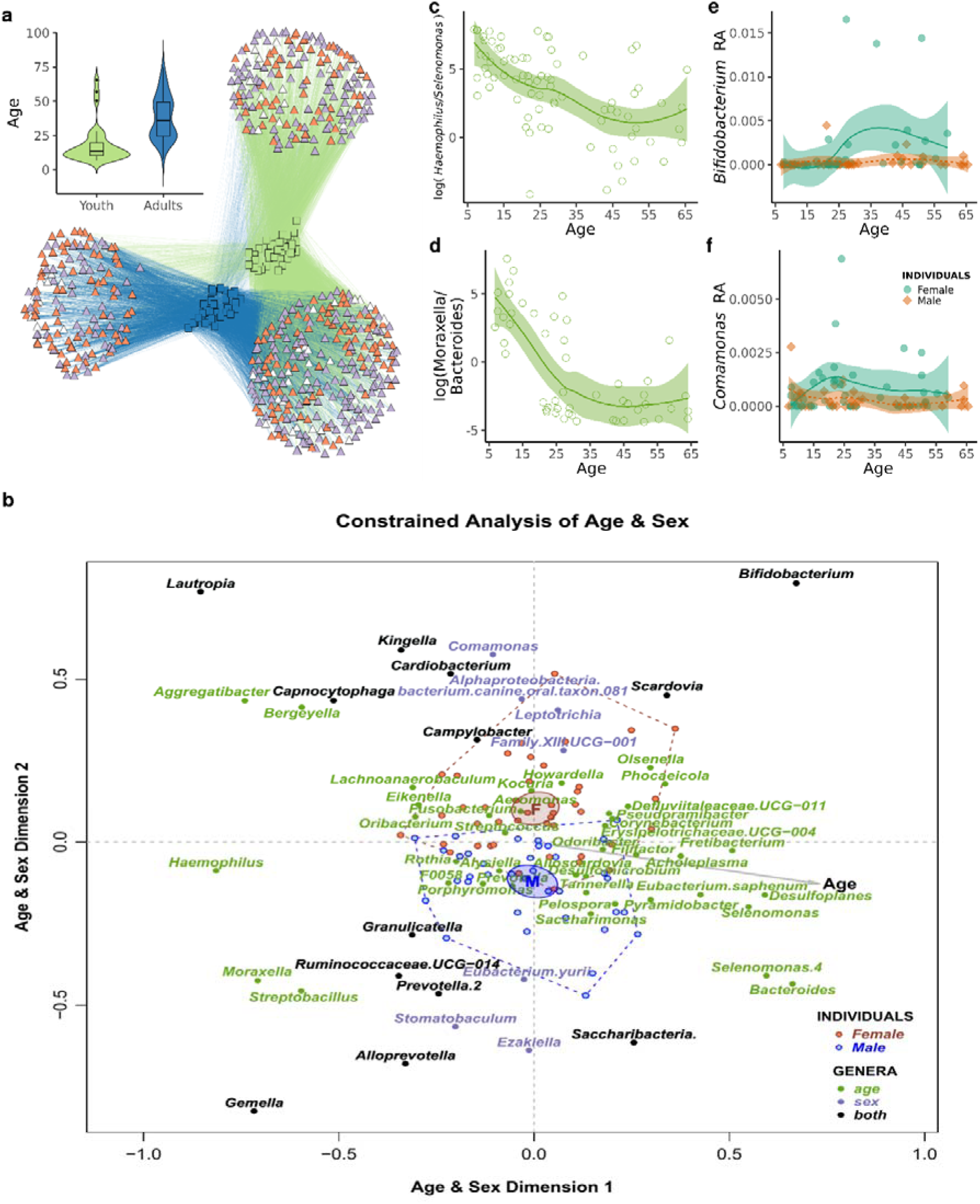
Age and sex-related effects in the hunter-gatherer oral microbiome. a) Network representation of the hunter-gatherer CMM. ASVs (triangles) are colour-coded as: putatively pathogenic (purple), non-pathogenic (orange) or unclassified (white). Inset shows age distribution for the two clusters of individuals (squares). b) Logratio analysis constrained to age and sex differences on the bacterial composition at genus level. The effects of diet were partialed out. Only genera statistically significant in at least 20 (for age) or 10 (for sex) logratios are displayed (p-value < 0.05 after Benjamini-Hochberg correction). Dashed lines enclose all individuals (dots) within a category of sex, with 95% confidence ellipses for their means. Taxa are colour-coded depending on the associated variable: age, sex, or both. The starting point of the grey arrow indicates the mean age of the population (30 years old). Log ratio of c) *Haemophilus* and *Selenomonas* abundance and d) *Moraxella* and *Bacteroides* according to age. Line and shaded area indicate the 95% confidence interval of the mean. Relative abundance of e) *Bifidobacterium* and f) *Comamonas* according to age and sex. Lines and shaded areas indicate the 95% confidence interval of the mean for each sex.

There is a clear change in the composition and frequency of certain bacteria with age (Figure 1c-d). At young age, we observe organisms that typically live in mucosa, such as *Haemophilus* and *Moraxella*, that infect the upper and lower respiratory tract but are detected in the oral cavity and saliva which are their vehicles of transmission. Other genera found at younger ages include bacteria normally associated to good oral health, such as *Bergeyella* and *Rothia*^14^. However, at older ages we observe a marked decline in the abundance of those genera and an increase of important pathogens related with periodontitis including the “red complex” periodontal pathogen *Tannerella*, as well as other periodontitis-related bacteria (*Filifactor, Fretibacterium, Saccharimonas, Selenomonas*, and *Phocaeicola*), consistent with a higher incidence of this disease with older age^15^. We also found organisms associated with cavities (*Olsenella*), with dental plaque and dental calculus formation (*Corynebacterium*), with pulmonary infections, sepsis, or bacteremia, and with chronic diseases (*Acholeplasma*) (see Methods for in-depth bacteria pathogenic classification). Another sign of ageing was the presence in the oral cavity of gut bacteria (*Bacteroides*) indicating a potential age-related decline in immunological function and filtering^16^. However, such changes are not associated with a decrease in the alpha-diversity of the total oral microbiome as measured by the number of bacteria observed or their phylogenetic complexity (Supplementary Figure 6a-b), suggesting that the overall effect of aging is a replacement of protective and commensal bacteria by pathogenic ones. This is supported by an increase in the number of potential pathogenic bacteria in the CCM in bacterial clusters associated with older ages (Fisher exact test, P < 0.001) (Figure 1a).

### Sex differences shape composition but not diversity of the Agta oral microbiome

We found no differences in alpha-diversity in the Agta oral microbiome between males and females (Supplementary Figure 6c-d), which may be explained by sex equality within the Agta hunter-gatherer society regarding diet and social interactions^17,18^. Nevertheless, the LRA constrained to age and sex shows sex-related differences in the composition of the oral microbiome (Figure 1b). For example, *Stomatobaculum* and *Eubacterium yurii*, present in the oral cavity of smokers^19^, are associated with males, consistent with Agta men chewing tobacco more frequently than women. It is also interesting to mention *Comamonas* (Figure 1f), which even if it has been reported as a possible contaminant in microbiome studies^20^, its presence in females could be related to its capacity of degrading the female hormone progesterone^21^. This bacterium has been found in subgingival samples, where female hormones could be present either in saliva or in gingival crevicular fluid.

### Age and sex interactions in microbiome composition

Some bacteria are significantly associated to both age and sex-related differences, such as *Gemella*, which is a prevalent inhabitant of the respiratory mucosa such as *Haemophilus* and *Moraxella*, supporting the idea that mucosa-associated and/or respiratory-tract organisms are more frequently acquired in young individuals, especially males. At older ages, the *Bifidobacterium*/*Saccharibacteria* ratio distinguishes between sexes: while *Bifidobacterium* is associated to females, the periodontal pathogen *Saccharibacteria* is associated to males. Thus, the observed trend of increase of periodontal pathogens with age is stronger in males, as expected by the global epidemiology of the disease^22,23^. On the other hand, we found an increase of the caries-related pathogens *Scardovia* and *Bifidobacterium* associated to reproductive age females. Caries incidence increases with age and is more prevalent in females^24^, a more saccharolytic or acidic salivary environment in older women, together with hormonal fluctuations and lower salivary flow^25^ could facilitate the proliferation of saccharolytic bacteria. The strong association of *Bifidobacterium* with adult females could also be explained by its presence in breastmilk^26^. Also, its proliferation coincides with the start of the reproductive age and the increase of childcare^10^ (Figure 1d).

### Influence of variation in rice consumption on the Agta oral microbiome

While the impact of diet on gut microbiome has been clearly established^27–30^, its role in the oral microbiome is still unclear. Some studies have found little or no effect, whereas others have found associations with specific nutrients^2,25,31,32^. The variation in rice consumption in the Agta allows us to assess both the relationship between the hunter-gatherer diet on the oral microbiome and the effects of the recent introduction of farming products. We performed a LRA on the bacterial genus abundance after partialing out the effects of age and gender (Figure 2). The first dimension of the ordination shows the gradient of the transition from a diet where all meals include meat to where most consist of only rice. Agta following a hunter-gatherer diet, where most meals contain meat, have large quantities of *Actinobacillus, Alphaproteobacteria* and *Streptobacillus* and lower abundance of *Selenomonas, Atopobium, Peptoanaerobacter*, and *Pyramidobacter*. The higher abundance of *Actinobacillus* in individuals ingesting a protein-rich diet is particularly interesting, given the extraordinary proteolytic potential of *A. actinomycetencomitans*, a well-known oral pathogen with destructive effects in the gingival tissue and in aggressive forms of periodontitis^33^. At the other extreme, in individuals with a rice-rich diet, there is an increase of the highly saccharolytic dental caries pathogen *Scardovia*, of *Treponema*, of gut organisms like *Butyrivibrio* and *Erysipelotrichaceae*, and of *Eggerthia*, a rare organism isolated from dental abscesses. We also ranked the ASV based on whether they are more present than expected in individuals with high or low proportion of meals with only rice or with meat/fish. We found that the scores associating each bacterial species with these two nutrients are negatively correlated (Spearman’s rho = -0.47, P < 0.0001). This fits with a general separation of oral microorganisms in saccharolytic (caries-related, acidogenic and acidophilic) and proteolytic (gum-disease and halitosis related, alkalophilic and NH^4^ generators), as suggested in a metabolome-based study^25^. Our results suggest that more settled Agta, which consume more rice, experience a decline in oral health, confirming a general pattern of health decline due to a Neolithic-like diet and a more farming-derived lifestyle^9,34,35^.

**Figure 2.**
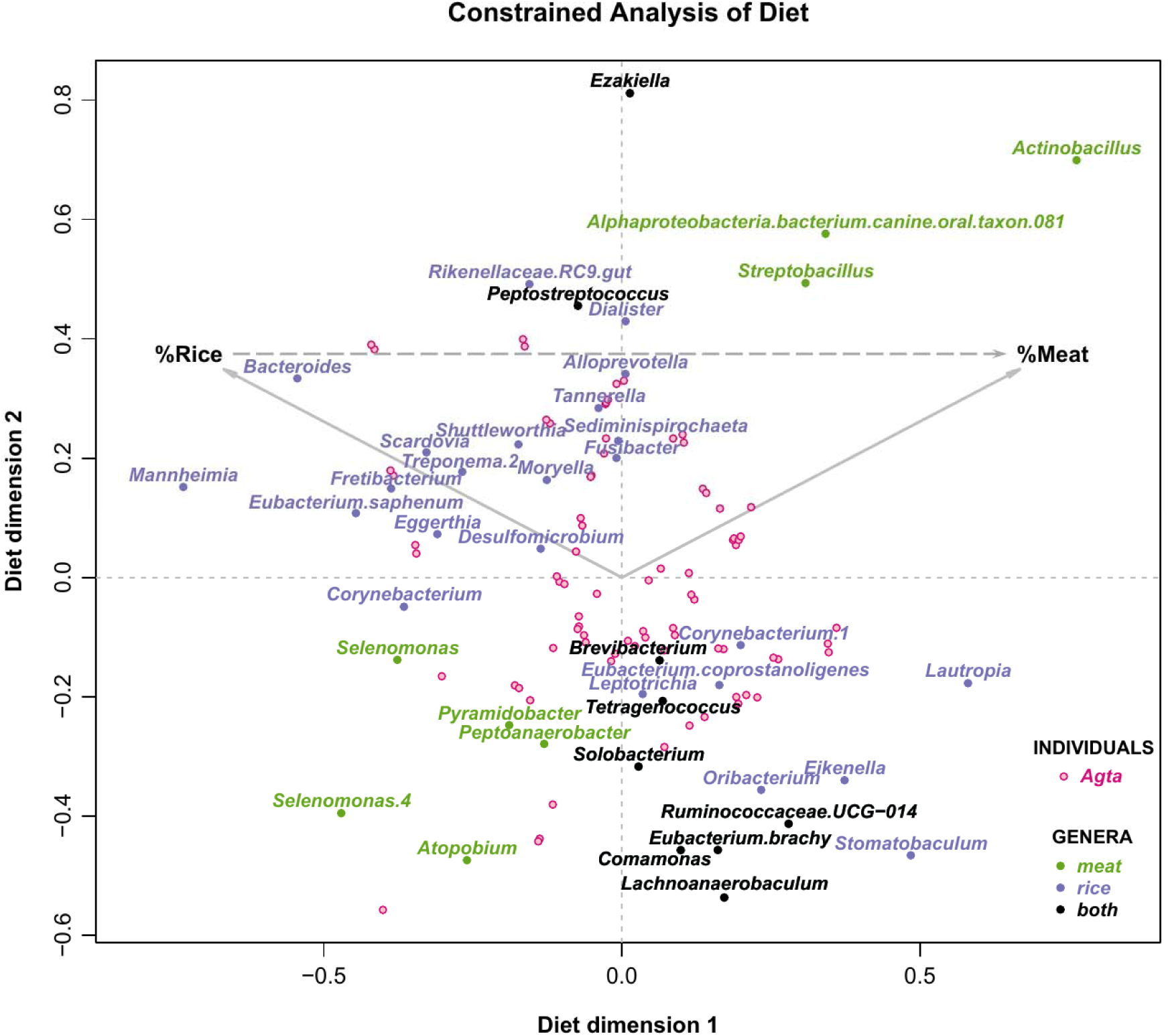
Effect of diet on the oral microbiome in the Agta. Logratio analysis constrained to diet differences on the bacterial composition at genus level. The effects of age and sex were partialed out. Only genera statistically significant in more than five (for rice) or three (for meat) logratios are displayed (p-value < 0.05 after Benjamini-Hochberg correction). Taxa are colour-coded based on the variable they are associated: proportion of meals with meat (%Meat), proportion of meals with only rice (%Rice), or both. The original result was slightly rotated so that the dashed vector indicating the difference between %Meat and %Rice was horizontal, without any change in explained variance.

### Pathogenic oral bacteria are associated with host collagen genes

The interaction between the host genetic makeup and microbiome composition differs across body sites^36,37^, and seems especially weak in the oral cavity^37,38^, making it difficult to assess the co-evolution of our genome and the oral microbiome. To overcome this, we performed a genome-wide association study (GWAS) using a mixed model approach in a population that evolved in a hunter-gatherer niche and without the confounding influence of antibiotics or brushing. We treated the relative abundance of each bacterium as an independent trait, adding age, sex, and household as covariates and kinship as a random effect. Household membership was used as proxy for the strength of social interactions between individuals, as social interactions predict microbiome sharing (Musciotto *et al. companion paper*). These analyses were performed using the CMM, and then using 92 genera present in at least 10 Agta. All bacteria identified in the Agta (Supplementary Table 1 and Supplementary Figure 7) overlap with those of other oral microbiome GWAS^3,36,37^ pointing to a small subset of oral bacteria influenced by the human genome (Figure 3). A pathway enrichment analysis linked this subset of the oral microbiome to several biological host functions (body fat metabolism, wound healing, and collagen trimmers) (Supplementary Table 2). Of relevance is an association between the pathogenic bacteria *Aggregatibacter* and *Selenomonas* with genetic variation in collagen genes. The ability to bind collagen is a vital feature in the oral cavity, as many oral bacteria require collagen-binding proteins to attach to oral tissues^39^ suggesting a genetic basis for the predisposition of biofilm formation by those bacteria. We further tested if we could detect signatures of positive selection in the host genomic regions associated with the oral microbiome, but we found no signals indicating recent selective pressures caused by oral bacteria.

**Figure 3.**
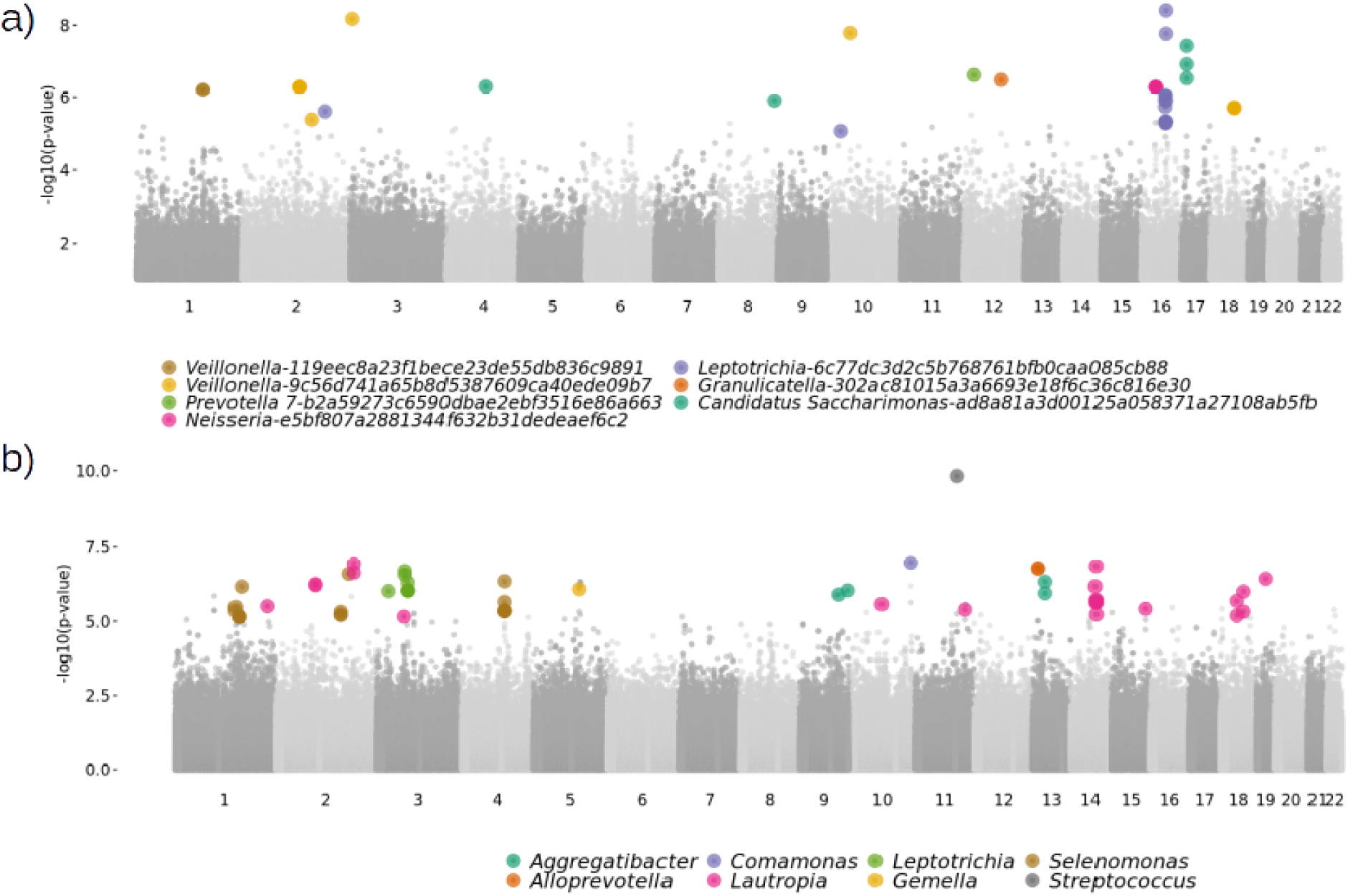
Genome-wide association study on bacteria abundance. Aggregated Manhattan plot of the GWAS results of the a) seven ASV and b) eight genera with non-zero PVE (“chip heritability”) estimates with at least one significant genetic association. Each dot is a SNP and significant SNPs-bacteria associations (q < 0.1) are color-coded according to the associated bacteria.

## Discussion

The Agta microbiome is influenced by external factors such as social interactions (Musciotto *et al. companion paper*) as well as intrinsic and ecological factors such as age, sex, diet and host genetics. Among the latter, we have shown that age has the strongest effect, with commensal or beneficial microbiota being replaced by potentially pathogenic ones with ageing. The proliferation of oral pathogenic bacteria exhibits sexual dimorphism, with caries-related (in females) and periodontitis-related (in males) bacteria increasing with age, likely associated with immunosenescence^40^ and with a sex-specific oral environment due to biological and cultural factors. In the Agta, the increase of farming-derived novel foods such as rice influences their microbiome composition and health. The relatively small subset of bacteria linked to the host genome, which are also found associated to other factors, suggests that the Agta oral microbiome is mainly affected by environmental (diet) and intrinsic factors (age), with little influence of individual host genetic variation (Supplementary Figure 5). Thus, environmental factors and not host genetics are the main driving force for oral microbiota acquisition, in agreement with Mukherjee *et. al*^41^. Based on the case study of the Agta hunter-gatherers, we conclude that the human oral microbiome is multifactorial with distinct subsets of bacteria shaped by specific ecological and social factors, reflecting multiple adaptations in the domains of life history, sociality, and diet.

## Methods

### Ethics approval

This study was approved by UCL Ethics Committee (UCL Ethics code 3086/003) and carried out with permission from local government and community members. Informed consent was obtained from all participants, after group and individual explanation of research objectives in the indigenous language. A small compensation (usually a thermal bottle or cooking utensils) was given to each participant. The National Commission for Indigenous Peoples (NCIP), advised us that the process of Free Prior Informed Consent with the tribal leaders, youth and elders would be necessary to validate our data collection under their supervision. It was done in 2017 with the presence of all tribal leathers, elders and youth representatives at the NCIP regional office, with the mediation of the regional officer and the NCIP Attorney. The validation process was approved unanimously by the tribal leaders, and the NCIP, and validated the full 5 years of data collection.

### Saliva sample collection

Saliva samples from 155 Palanan Agta were collected over two field seasons: April-June 2013 and February-October 2014. For comparative genetic studies we also used saliva samples from 21 Mbendjele Bayaka, an African hunter-gather population, collected in 2014, and 14 Palanan farmers collected in 2007-2009. In all cases saliva was collected using the Oragene·DNA/saliva kit and participants were asked to rinse their mouth with water and to spit into the vial until half full. After collection and transportation, saliva samples were stored at the UCL Department of Anthropology, London, UK at -20ºC.

### Microbial DNA extraction and 16S rRNA gene sequencing

A total of 155 Agta saliva samples were selected to study their microbiome composition. DNA was extracted following the protocol for manual purification of DNA for Oragene·DNA/saliva samples. The 16S rRNA gene V3-V4 region was amplified by PCR with primers containing Illumina adapter overhang nucleotide sequences. All PCR products were validated through an agarose gel and purified with magnetic beads. Index PCR was then performed to create the final library which was also validated through an agarose gel. All samples were pooled together at equimolar proportions and the final pool was qPCR quantified prior to the MiSeq loading. Raw Illumina pair-end sequence data were demultiplexed, quality filtered and denoised with QIIME 2 2019.1^42^ and DADA2^43^. DADA2 generates single nucleotide exact amplicon sequence variants (ASVs). ASV are biological meaningful entities as they identify a specific DNA sequence and allow for higher resolution than using operational taxonomic units (OTUs)^11^. Taxonomic information was assigned to ASVs using a naïve Bayes taxonomy classifier against SILVA database release 132 with a 99% identity sequence^44^. ASVs that did not belong to the kingdom Bacteria, or that were classified as mitochondrial or chloroplast sequences and samples with an extremely low number of sequences (8000) were excluded from further analyses. ASVs were aligned with MAFFT^45^ and a rooted phylogenetic tree was constructed with FastTree2^46^ using default settings via QIIME 2. This resulted in a total of 5430 ASVs and 138 Agta (67 women and 71 men). We generated a rarefaction curve with R package vegan (version 2.5-7)^47^ to determine that the richness of the samples had been fully observed (Supplementary Figure 8). The number of observed ASVs and Shannon Diversity index were calculated with R package Phyloseq (version 1.30.0)^48^. Faith’s Phylogenetic Diversity index^49^ was calculated with R package picante (version 1.8.2)^50^ using the rooted phylogenetic tree generated in R^51^. To determine the set of microbial traits to be included in the analyses, we selected ASVs with at least 10 reads in at least 2 individuals (n = 1980), then we aggregated those with a taxonomic assignment at a genus level, resulting in 110 genera. At the ASV level we also defined a Core Measurable Microbiota (CMM), consisting of ASVs that appear in at least 10% of the Agta individuals (14 or more) resulting in 575 ASVs (out of 1980) that represent 90% of the composition of the Agta microbiome.

### Genotype data

A total of 190 saliva samples were genotyped with the Affymetrix Axiom Genome-Wide Human Origins 1 array. DNA extraction was carried out following the protocol for manual purification of DNA for Oragene·DNA/saliva samples in the same laboratory that sequenced the 16S rRNA data. Samples were analyzed with Axiom Analysis Suite v4.0 following the Axiom genotyping best-practices workflow for saliva samples. 618810 markers and 177 samples passed initial quality control. Single nucleotide polymorphisms (SNPs) with less than 95% genotyping rate and samples where the estimated gender from the genotypes did not match the recorded gender were excluded from the analysis. Duplicated samples were identified with KING^52^ and removed. This resulted in a total of 617063 markers and 174 samples: 141 Agtas, 19 Bayaka and 14 Palanan farmers.

### Ethnographic data collection

Ethnographic data collection occurred over two field seasons from April-June 2013 and February-October 2014. In the first season we censused 915 Agta individuals (54.7% which were male) across 20 camps, collecting basic information on household composition, sex, and estimated ages. Following relative aging protocols^53^, accurate ages were established for all individuals post data collection.

### Diet data collection

Dietary recall data was collected at the household level over a period of 10 days. We asked the mother and the father at the end of the day (between 17:00 – 18:00) what foods they had eaten that day, including agricultural produces from trade with nearby farmers. We counted the total amount of meals we had recorded for a household and established what proportion of these consisted of meat, vegetables, fruits, honey, and rice. Therefore, this is only a rough guide to dietary composition and does not take calorific intake or absolute weighs of the different food types into account. Dietary data for 80 individuals (37 males and 43 females) was annotated based on the proportion of meals that consisted of only rice, and the proportion of meals that included meat (primarily fish and other marine resources and game).

### Classification of oral bacteria as pathogens

Bacteria were classified as potential oral pathogens if they have been reported as etiological agents of periodontitis or dental caries. Assignment as periodontal pathogen was performed according to the systematic review of Perez-Chaparro *et al*.^54^ and Socransky *et al*.^55^, or if they have been previously associated with this gum disease^56,57^. Bacteria were classified as caries pathogens if they were described in transcriptomic studies of human cavities, according to Simon-Soro *et al*.^58^ and Simon-Soro and Mira^59^, previously associated with caries^60,61^, with cavities^62^ or with dental plaque and dental calculus formation^63,64^. Bacteria reported as etiological agents of respiratory infections and biofilm-mediated infections were also considered pathogens, including organisms that can be present in healthy carriers. These included species described in Leung *et al*.^65^, Bellussi *et al*.^66^, and Natsis and Cohen^67^. Bacteria causing urinary tract infection or sexually transmitted diseases which can transiently be found in the oral cavity were also considered as potential pathogens and included microorganisms described in Lanao *et al*.^68^ and Jung *et al*.^69^. Common oral commensals potentially causing endocarditis or systemic infections in immunocompromised patients were not considered pathogens. If a bacterium was isolated from the oral cavity of an animal, it was considered an oral inhabitant for the sake of our classification. If taxonomic classification in our dataset could be assigned at the genus level only, it was considered a pathogen if: i) > 90 of species within the genus were pathogenic, or ii) it included a major pathogenic species but the rest of species within the genus were not oral inhabitants, according to the Human Oral Microbiome Database (http://www.ehomd.org/)^70^. Bacteria with taxonomic assignments at higher levels than genus (family, order, class) were excluded from this analysis.

For assignment of bacteria to pathogenic or non-pathogenic, we used species-level ASVs, given that there are multiple cases where different species from the same genus had a different assignment. If taxonomic classification of the ASV was only possible at the genus level, it was considered a pathogen if: i) >90% of named species within the genus were pathogenic, or ii) the genus included a major pathogenic species but the remaining species within the genus were not classified as oral by the Human Oral Microbiome Database^70^. ASV with a top hit to a sequence classified as “Oral taxa” in databases but without a species assignment were not considered named species and were discarded from the analysis. Cases where taxonomic classification of the ASV was only possible at the family level or higher were also discarded.

### Multivariate compositional data analysis on microbial composition

We performed a constrained logratio analysis (LRA) using the package easyCODA^12^ in R^51^ on the Agta oral microbiome at the genus level using as constraining covariates age (as continuous variable), sex (male and female), and diet (both proportion of meals with meat and proportion of meals with only rice). The microbiome abundance counts of each Agta individual were treated as compositional data^71^ and transformed to logarithms of ratios (logratios). Constrained LRA is a special case of redundancy analysis^12^ where the total logratio variance is decomposed into parts explained by the covariates (the “constrained variance”) and a residual part (the “unconstrained variance”, unrelated to the covariates). Then, the ordination resulting from the LRA explains a maximum of the constrained variance in a reduced two-dimensional solution. The statistical significance of the three covariates was assessed using a multivariate permutation test (999 permutations) in the R package vegan^47^. There is no correlation between these three covariates, except within the diet covariate, where the two variables are negatively correlated (Spearman’s rho = -0.54, P < 0.0001) (Supplementary Figure 1). To focus on the genera affected only by internal factors (age and gender), we performed a constrained LRA on the microbial composition after partialing out the effects of diet.

Similarly, to identify genera affected exclusively by the diet, we performed a constrained LRA after partialing out the effects of age and gender. Taxon-covariate association was ranked by counting the number of significant logratios for each of the taxa, with p-value < 0.05 controlling for the false discovery rate (FDR) at level _α_ = 0.05^72,73^.

### Community detection

To model the relationship between the Agta and the CMM, we used a stochastic block model (SBM) approach specifically suited for bipartite networks^13^. SBM infers the community structure^74^ that better fits the existing graph, by building a prior distribution for edges that holds no information on real data and using it in the framework of Bayesian inference (biSBM) to find a partition of the two types of nodes whose associated entropy is maximal. In this framework, the absence of links between nodes of the same type or set is not considered informative for the model, as it is expected given the bipartite nature of the graph, different from the general version of SBM. We selected the number of clusters in the two sets that minimize the description length^75^. Robustness of the clustering was assessed by calculating the average Adjusted Rand Index (ARI) between iterations (n = 100), finding a mean ARI on Agta = 0.90 and a mean ARI on ASV = 0.70. ARI measures the similarity of two partitions against a null hypothesis of random assignment maintaining the size of the different clusters; the closer to 1 the more robust is the classification^76^. The resulting clusters were plotted with graph-tool^77^.

### Ranking of bacteria associated with diet

ASVs present in the Agta were ranked from -1 to 1 based on whether that ASV is more present than expected in individuals from a given category: low proportion versus high proportion of meals with only rice, and low proportion versus high proportion of meals with meat, based on the median value of the population for each variable. Thus, a meat associated score towards 1 indicates that an ASV is present more than expected in individuals with high proportion of meals with meat (above the median of the population), and a score towards -1 indicates that is present more in individuals with low proportion of meals with meat (below the median of the population). A rice associated score towards 1 indicates that an ASV is present more than expected in individuals with high proportion of meals that consist of only rice, and a score towards -1 indicates that is present more in individuals with low proportion of meals with only rice.

### Microbiome genome-wide association studies

To study the relationship between host genetics and the microbiome in the Agta, we used a genome-wide association study (GWAS) approach to identify specific SNPs associated with microbial abundance using GEMMA (version 0.94)^78^. GWAS were performed using the relative abundance of a given taxon in the Agta as a phenotype trait, adding as covariates age, sex, and household (as a proxy for diet and shared environment, as members of the same household share a hearth and their food on daily bases). A kinship matrix calculated by KING using identical by descent segment inference^52^ was included as random effects. For the GWAS analyses, we applied the following quality control steps to the Agta genotypes. First, to detect ancestry outliers in the dataset, we filtered the samples to keep only bi-allelic autosomal SNPs with Minor Allele Frequency (MAF) > 5% and without missing data with PLINK 1.9^79^. This dataset was pruned for linkage disequilibrium (LD) using --indep-pairwise 50 5 0.2, and we performed a principal component analysis (PCA) with EIGENSOFT (version 7.2.1)^80^ to identify ancestry outliers and exclude them from the analysis (Supplementary Figure 9). Second, per sample heterozygosity was calculated with PLINK and samples with overall increased/decreased heterozygosity rates (±3 s.d. from the mean of the population) were removed. A total of 129 Agta samples passed microbiome and genotype data quality controls and were included in the microbiome GWAS analyses.

The analyses were done at the genus taxonomic level and on the CMM to study the effect of host genetics at different taxonomic levels. Depending on the taxon analyzed, as we included only samples with non-zero abundance, and SNPs with MAF < 10% and with more than 5% missing data were excluded, the number of individuals tested ranged from 10 to 129 and the number of SNPs tested ranged from 270569 to 313198 markers. When we performed the GWAS at the genus level, we only included in the analysis 92 genera that are present in at least 10 Agta individuals to exclude low prevalent genera. P-values were adjusted for multiple testing by FDR, and SNP-taxa associations were considered significant at q-value < 0.1 on the cases where the proportion of variance in the bacterial abundance explained by the genotypes (PVE or “chip heritability”) was non-zero. The proportion of variance in the phenotype (bacterial abundance) explained by the genotypes tested (PVE or “chip heritability”) was estimated for each taxon and was considered non-zero if the standard error measurements did not intersect zero. We applied genomic control to correct for cryptic relatedness and population stratification and minimize false positives induced by inflated association test statistics^81^. To do so, we estimated the genomic inflation factor as the median value of the likelihood ratio test (LRT) values divided by 0.456 (median of a χ^2^(1) distribution) and recalculated the p-values after dividing the LRT values by the genomic inflation factor^82^. Threshold of significance was set at FDR 10%, and only genomic positions having at least three samples for the major homozygous genotype and for the heterozygous genotype were considered. SNPs were annotated with ANNOVAR^83^ in GRCh37 (hg19) using RefSeqGene and dbSNP 147. For the enrichment analyses, we extracted the genes associated to all non-intergenic SNPs and classified the genes in the background set, that consisted in all genes present in the Axiom Human origin array; and the set to test, that consisted of all genes that had a non-intergenic SNP significantly associated with an ASV or a genus. We performed a gene ontology enrichment analysis with ViSEAGO^84^ and TopGo^85^ R packages (Fisher exact test) and FUMA GENE2FUNCTION module (Functional Mapping and Annotation of Genome-Wide Association Studies)^86^ to perform pathway enrichment analysis (hypergeometric test) with FDR 5%.

### Selection analyses

To test whether the GWAS SNPs showed any signal of recent positive selection we performed a genome-wide scan of selection. We phased the Agta and Palanan farmer populations independently. For each population, samples that were identified as ancestry outliers by a PCA or with overall increased/decreased heterozygosity rates (±3 s.d. from the mean of the population) were excluded from phasing. A total of 138 Agta samples were phased using SHAPEIT2 version v2 (r900)^87^ with the duoHMM method to improve the phasing by integrating the known pedigree information. SNPs with missing data were removed and window size was set to 5Mb for phasing. Due to the small sample size, to phase the 14 unrelated Palanan farmers we used SHAPEIT2 with default parameters and the 1,000 Genomes Phase 3 panel of haplotypes^88^ as a reference dataset. SNPs with missing data were removed. For the selection analyses, we excluded one of each pair of related individuals by removing the sample in the pair with the lowest call rate in the Agta phased dataset. This resulted in 38 unrelated Agta individuals and 14 unrelated Palanan farmers. We ran the Integrated Haplotype Score (iHS)^89^ in the Agta phased dataset and the Cross-population Extended Haplotype Homozygosity (XP-EHH) test^90^ comparing the Agta against Palanan farmers as implemented in selscan version v1.3.0^91^ to identify signals of positive selection in GWAS SNPs. Both tests were run with default parameters and with the genetic map provided by the 1,000 Genomes Phase 3^88^. To identify regions under selection, for each test we selected markers with scores in the 95th percentile that had at least 3 markers in the 99th percentile in the surrounding area (± 10 Kb). For iHS we used absolute values, while only positive scores were analyzed for XP-EHH.

## Acknowledgements

A.B.M was funded by Leverhulme Trust Grant RP2011-R 045 and PLP-2017-323. J.B. received grant PID2019-110933GB-I00/AEI/10.13039/501100011033 from the Agencia Estatal Investigación (AEI), Spain, grant GRC 2017 SGR 702 from Secretaria d’Universitats i Recerca del Departament d’Economia i Coneixement de la Generalitat de Catalunya, as well as grant CEX2018-000792-M, part of the “Unidad de Excelencia María de Maeztu” funded by the AEI, which granted the microbiome analysis by the Servei de Genòmica at Universitat Pompeu Fabra. A.M. is funded by grant RTI2018-102032-B-100 from AEI (Spain). A.E.P. is funded by grant MR/P014216/1 from the Medical Research Council.

## Data availability

16S amplicon data (EGAS00001005317) are deposited at the European Genome-phenome Archive (EGA), which is hosted at the EBI and the CRG. Genome data generated in this study has been deposited at EGA under accession number EGAS00001005315. Data at the individual level on age, household composition and diet that support the findings of this study are available on request from the corresponding authors (JB and ABM). The individual data are not publicly available due to them containing information that could compromise research participant privacy.

## Code availability

Source code and data for visualization are available at https://doi.org/10.5281/zenodo.6342212

