## Supplementary Figures for "The making of the oral microbiome in Agta hunter-gatherers"

Begoña Dobon, Federico Musciotto, Alex Mira, Michael Greenacre, Abigail E. Page, Mark Dyble, Daniel Smith, Sylvain Viguier, Rodolph Schlaepfer, Gabriela Aguileta, Leonora H. Astete, Marilyn Ngales, Vito Latora, Federico Battiston, Lucio Vinicius, Andrea B. Migliano, Jaume Bertranpetit

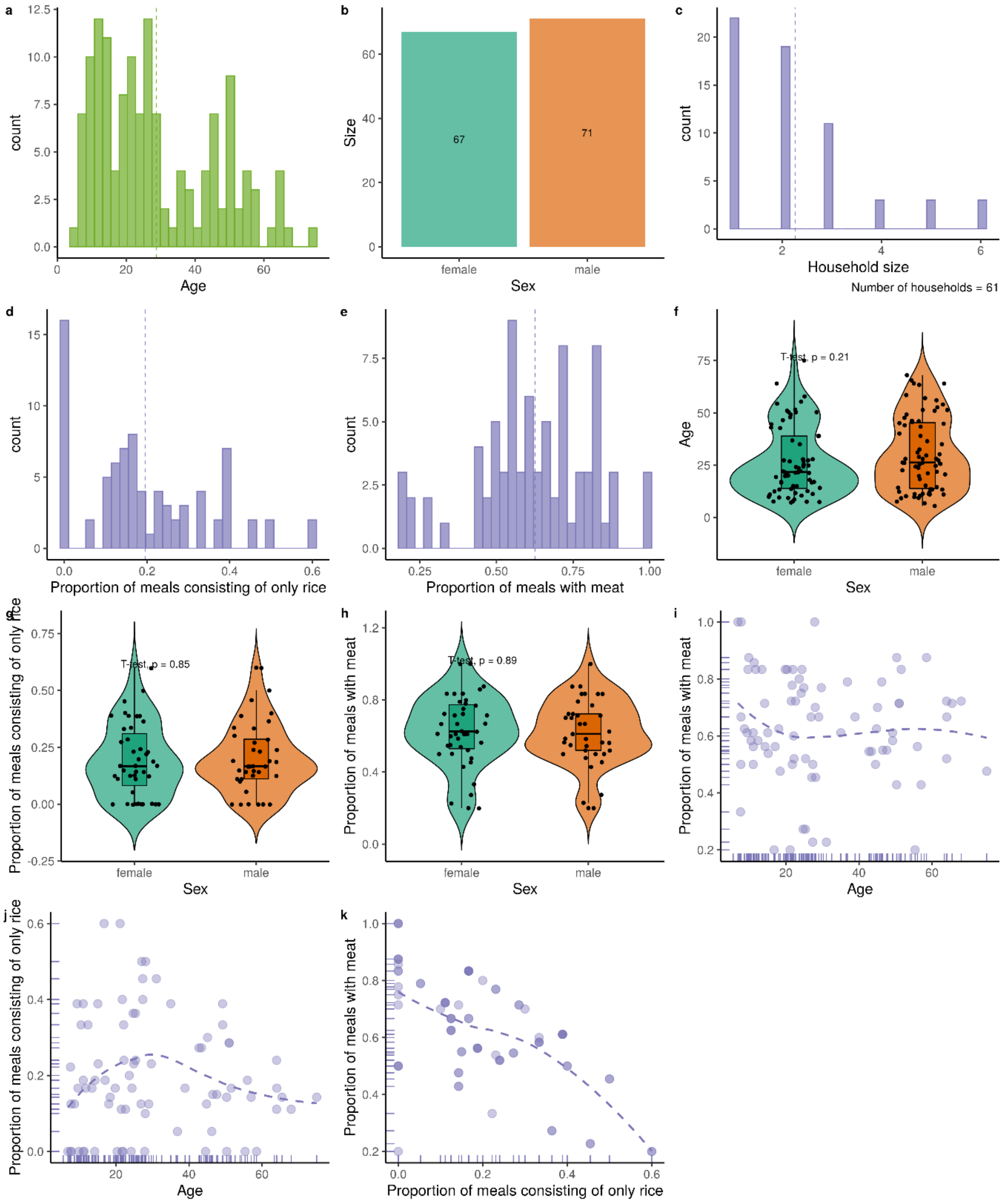

**Supplementary Figure 1. Distribution of the factors in the Agta population.** a) Distribution of age in the population. b) Number of individuals of each sex. c) Distribution of household sizes. d) Distribution of the proportion of meals with only rice. e) Distribution of the proportion of meals with meat. f) Distribution of ages according to the sex. g) Proportion of meals with only rice depending on the sex. h) Proportion of meals with meat depending on the sex. i) Proportion of meals with meat according to age. j) Proportion of meals with only rice according to age. k) Proportion of meals with only rice depending on the proportion of meals with only rice. Dashed line indicates the mean. Differences between sex categories were assessed by Student's t-test.

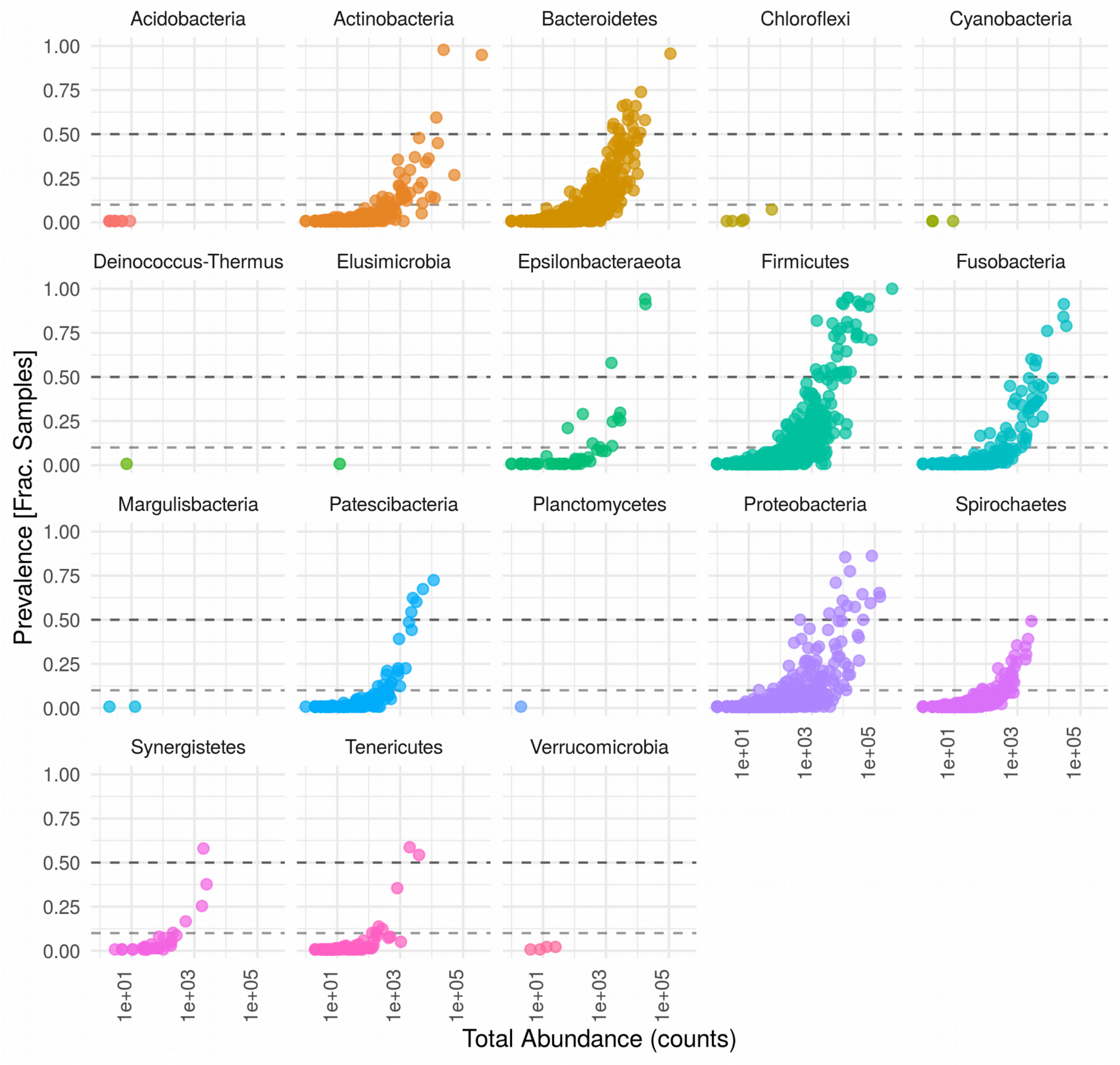


**Supplementary Figure 2. Phylum prevalence in the Agta oral microbiome.** Fraction of the samples where any of the ASVs is present at least once (prevalence) versus the total abundance of each phylum (number of counts). Horizontal dashed lines indicate when an ASV is present in 10% or 50% of the population.

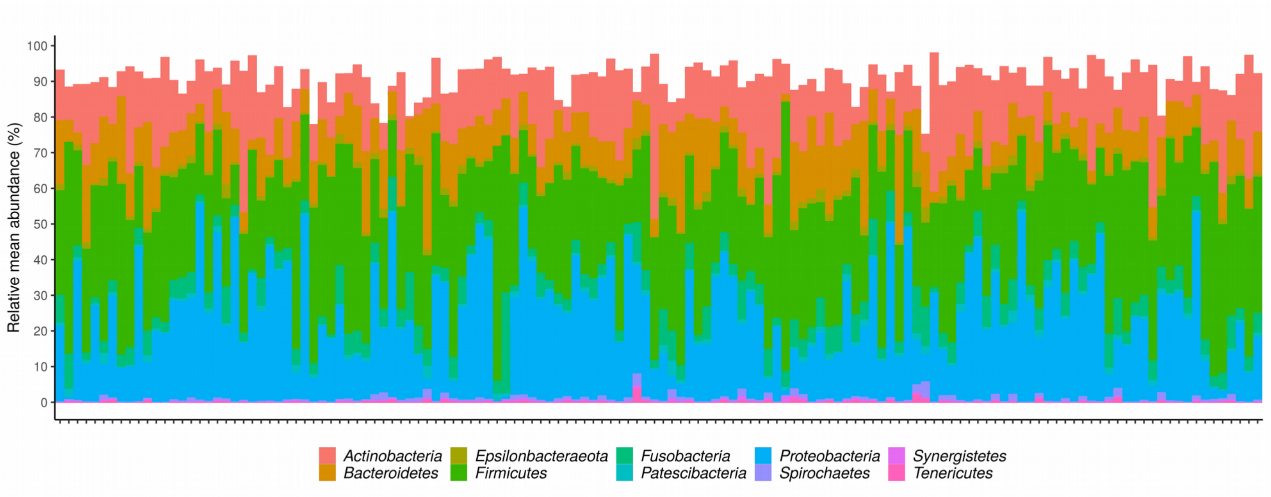


**Supplementary Figure 3. Core Measurable Microbiome.** Relative abundance of the 575 ASV that are part of the CMM aggregated at the phylum level for each Agta individual. The relative mean abundance for does not reach 100% as we only show ASVs belonging to the CMM.

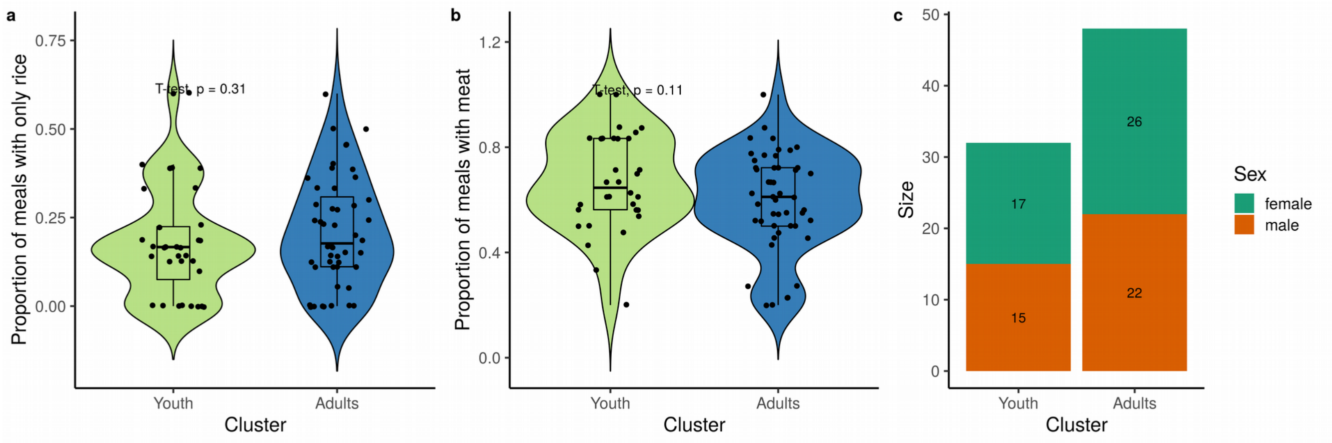


**Supplementary Figure 4. Distribution of the factors in the CMM age-based clusters.** a) Proportion of meals with only rice according to the age-based cluster. b) Proportion of meals with meat according to the age-based cluster. c) Proportion of males and females according to the age-based cluster. Differences between categories were assessed by Student's t-test.

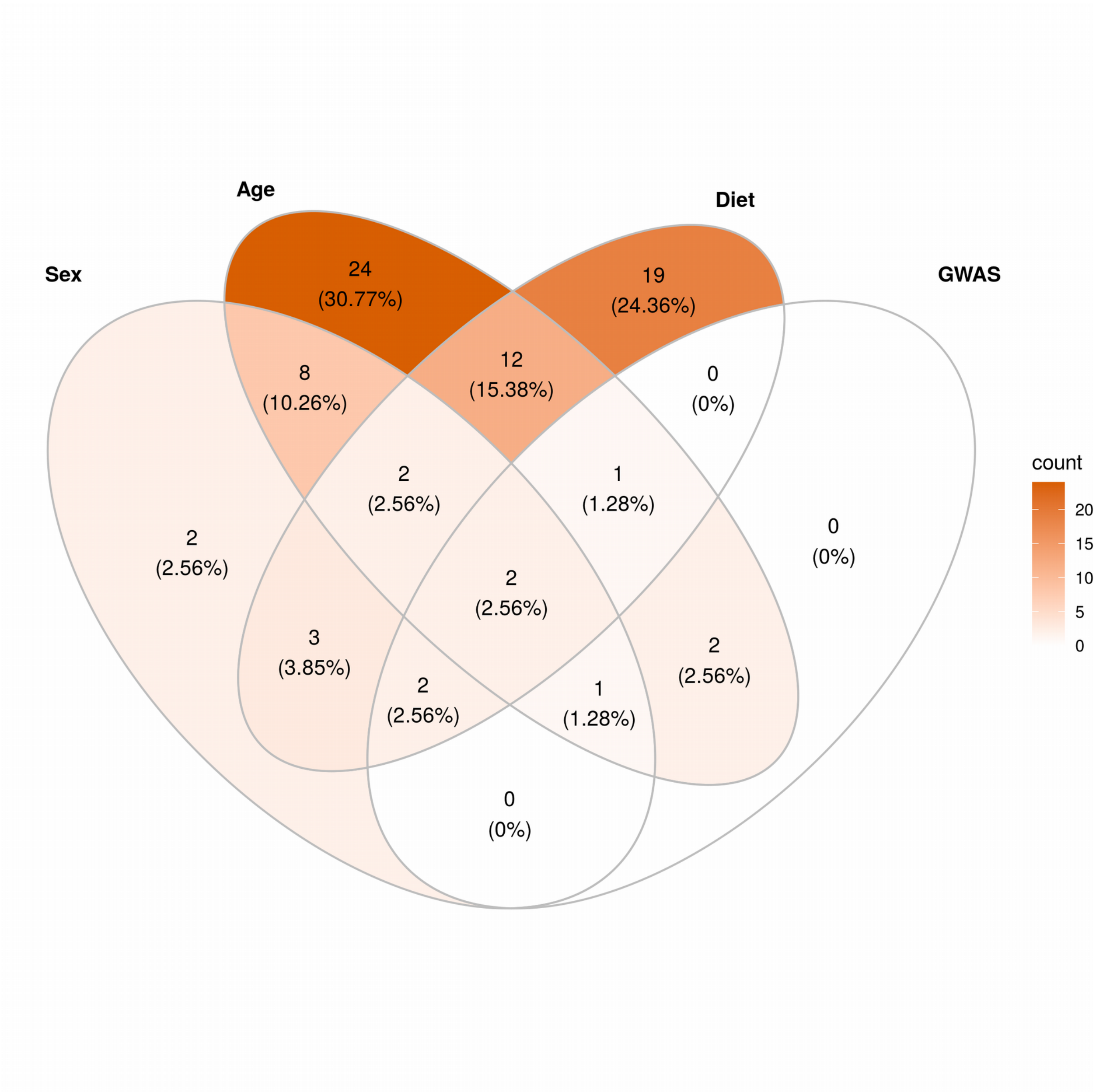


**Supplementary Figure 5. Intersection of the genera associated to each factor.** Venn diagram showing the intersection (number and percentage) between the genera significantly associated to each factor.

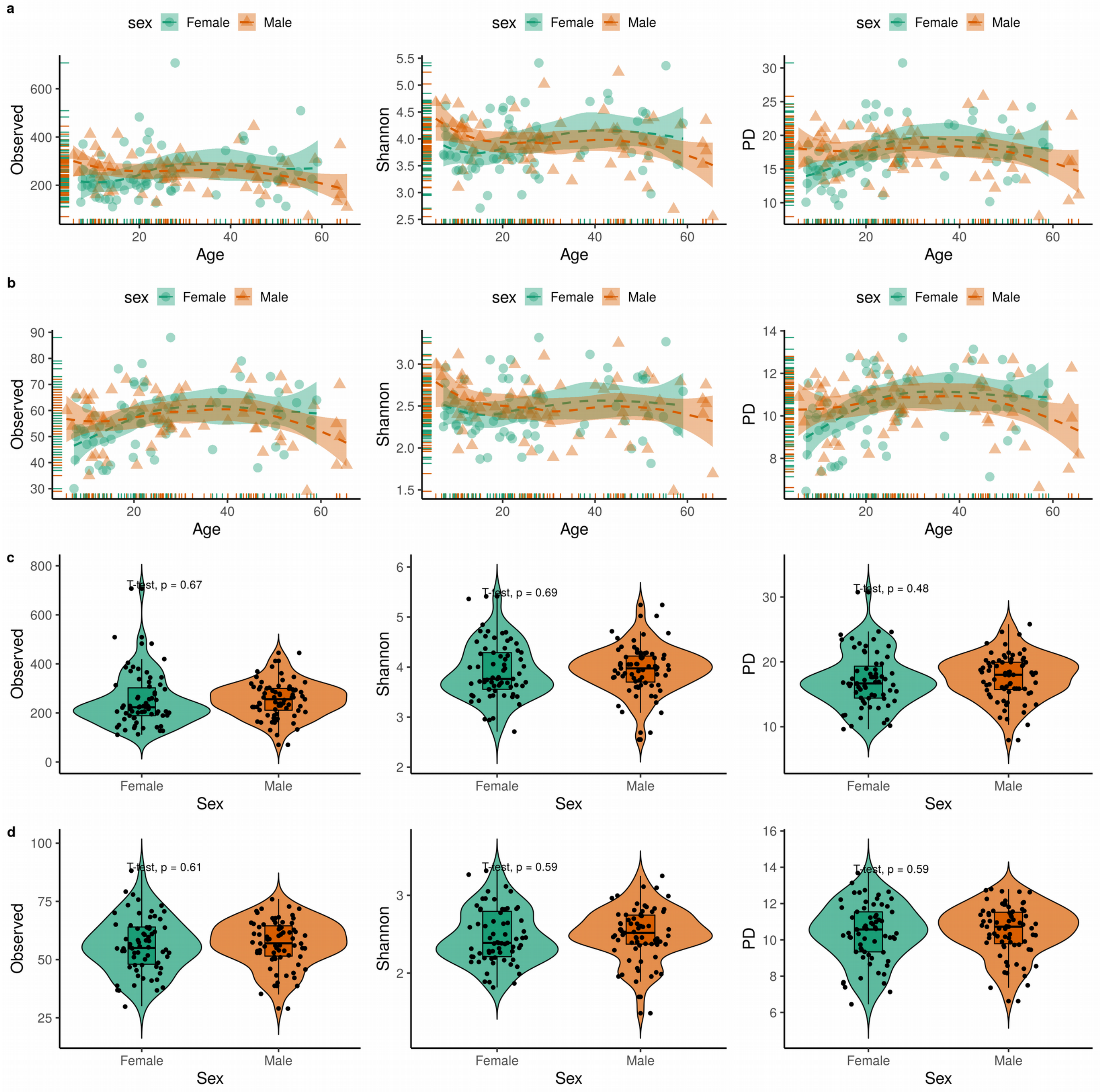


**Supplementary Figure 6. Alpha diversity in the oral microbiome.** a) Number of observed ASVs, Shannon diversity and Faith Phylogenetic Diversity index (PD) in each sample according to age and sex. b) Number of observed genera, Shannon diversity and Faith Phylogenetic Diversity index (PD) in each sample according to age and sex. c) Number of observed ASVs, Shannon diversity and Faith Phylogenetic Diversity index (PD) depending on the sex. d) Number of observed genera, Shannon diversity and Faith Phylogenetic Diversity index (PD) in each sample according to age. Dashed line and shaded area indicate the mean and the standard error. Differences between sex categories were assessed by Student’s t-test.

**a**

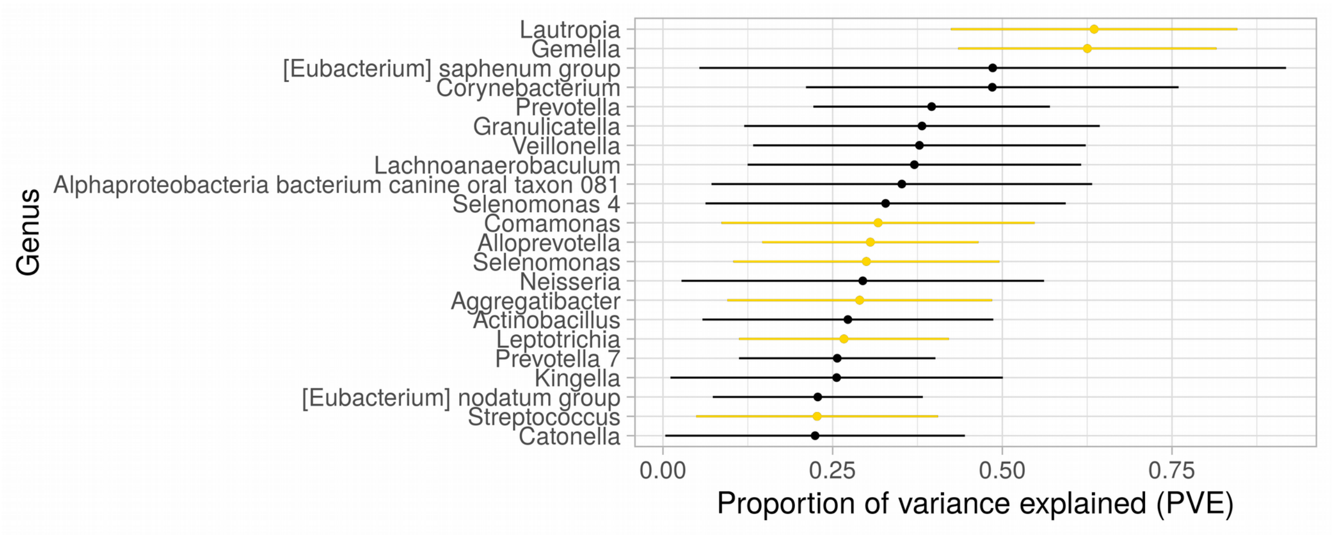


**b**

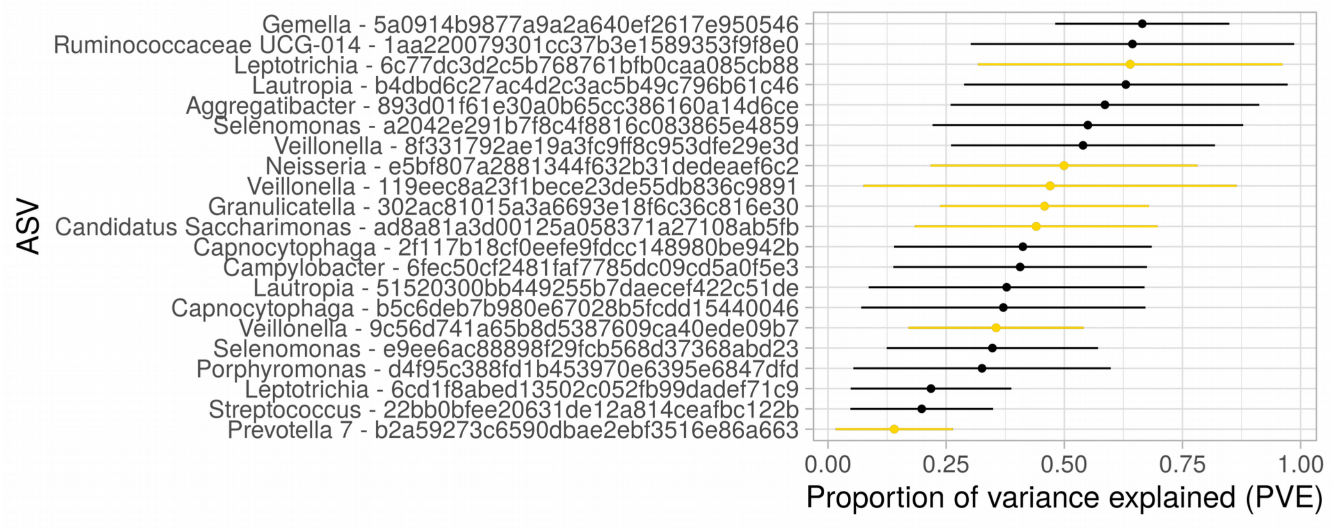


**Supplementary Figure 7.** Chip heritability for the Agta oral bacteria. Each point represents the estimated proportion of variance in bacterial abundance explained by all Agta genotypes analyzed (PVE, or “chip heritability”) in the microbiome abundance GWAS at the a) genus or b) ASV level (CMM). For the ASV is shown the ASV ID and the lowest taxonomic rank assigned. Bars represent the standard error of the PVE estimate. Only taxa with non-zero PVE estimates are shown. Bacterial taxa that have at least one significant genetic association at FDR 10% are highlighted in orange.

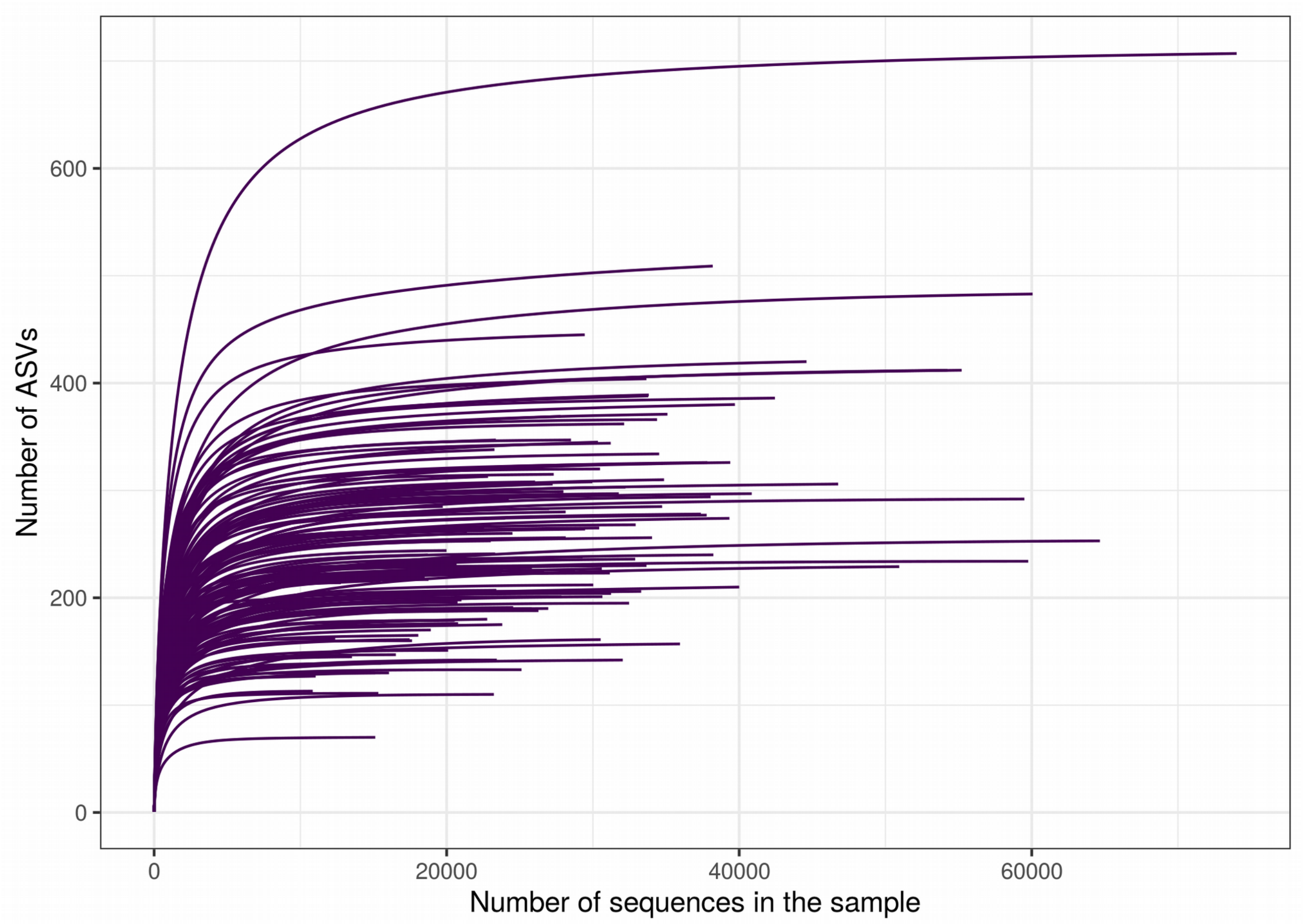


**Supplementary Figure 8. Rarefaction curves.** Rarefaction curves calculated at an interval step of 50 for each sample. Each line shows the cumulative number of different ASVs found in a sample based on the number of sequences sampled.

**a)**

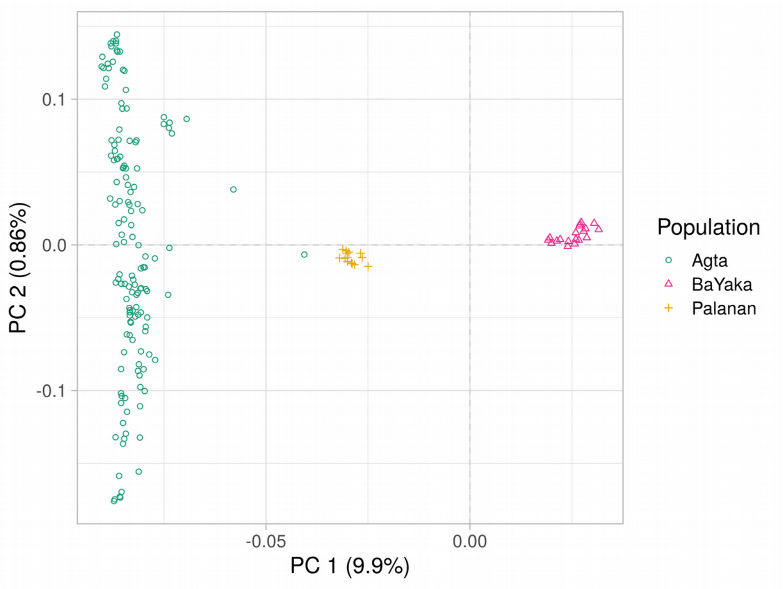


**b)**

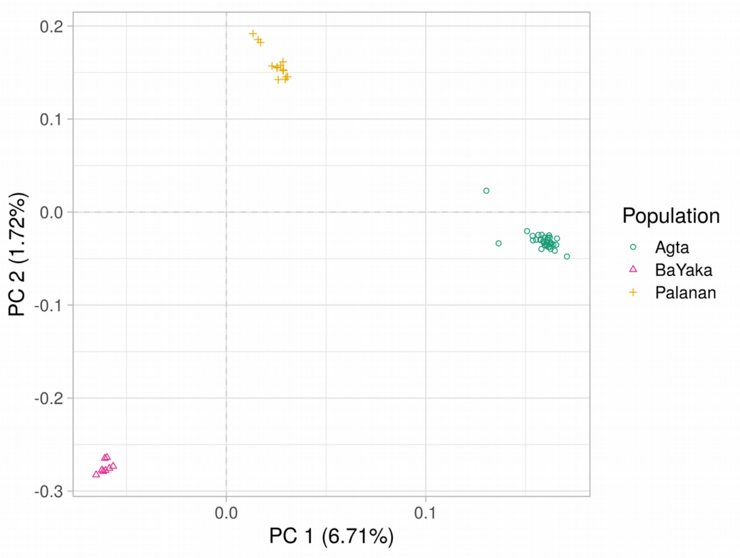


**Supplementary Figure 9.** Principal component analysis. Principal component analysis of the three populations genotyped. a) First two principal components (PC) showing the Agta, the Bayaka (African hunter-gatherer) and the Palanan farmers. A few Agta individuals are closer to the Palanan farmer population than to other Agta. b) PC1 and PC2 after removing ancestry outliers, closely related individuals, and samples with extreme heterozygosity levels.
